# Amplified transmission in host communities following disease-induced declines

**DOI:** 10.64898/2026.08.18.744677

**Authors:** Joseph R. Hoyt, John E. DePue, Macy J. Kailing, Nichole A. Laggan, Heather M. Kaarakka, Jennifer A. Redell, J. Paul White, Kate E. Langwig

## Abstract

Pathogen transmission is a fundamental component of infectious disease systems, governing the speed and extent of pathogen outbreaks. Predicting transmission within and among species is therefore critical for outbreak preparedness and the effective implementation of control strategies. Epidemics themselves can alter host populations in ways that fundamentally reshape transmission, but the long-term consequences remain poorly understood. Here, we quantify changes in transmission using a surrogate pathogen in bat communities before and after the emergence of white-nose syndrome, a fungal disease that has caused widespread population declines across North America. We find that transmission increased following disease-induced declines across all species in the community. We also observed an increase in environmental contacts and expanded habitat use by individuals, suggesting that surviving bats, on average, increased their activity and opportunities for interaction despite the substantial reductions in density and costs of infection. Together, these findings demonstrate that pathogen emergence can fundamentally alter host contact patterns and transmission dynamics, emphasizing the importance of considering behavioral and ecological changes imposed by epidemics when predicting pathogen spread in naïve host populations.

## Introduction

Transmission among hosts is a critical part in the life history of all pathogens^1–5^. The rate at which new individuals become infected determines the speed and extent of an epidemic, the breadth of host population impacts^4,6–9^, and can influence whether certain intervention and control measures are successful^4,10,11^. Predicting how a pathogen might move through a population or community of hosts can allow rapid response upon pathogen introduction and more targeted control efforts^6,12–14^. However, transmission is a dynamic process shaped by interactions among hosts, pathogens, and their environment, and there are numerous factors that can influence the rate of contacts and transmission over time^15–18^.

Processes inherent to pathogen introduction events are predicted to directly influence the pathways and magnitude of transmission^12,19–21^. These include changes in host abundance^22–24^, behavior^3,25–28^, community composition^29–34^, pathogen evolution^35–38^ and host adaption (resistance)^39–41^. Reductions in host density following disease mortality events, which often accompanies novel pathogen introduction, can directly reduce transmission for pathogens that are density dependent ^22,42–44^. For pathogens that cause high mortality or sterility, behaviors and interactions that increase exposure risk should also impose substantial fitness costs to the host^25,45–50^. However, given that these behaviors likely conferred fitness benefits prior to pathogen introduction^3,51,52^, it remains unclear whether hosts will reduce interactions to limit transmission or maintain high contact behaviors despite the increased risk of infection^53,54^. In systems where transmission remains high, hosts will need to adapt other strategies to reduce disease^23,55–60^. However, it can be difficult to disentangle behaviorally driven changes in transmission from other mechanisms that alter transmission over time^1,61,62^, as these processes often occur simultaneously following pathogen invasion. In addition, empirically estimating transmission is a challenge across most disease systems^61,63–67^, which has limited our ability to understand and predict how or why transmission changes following pathogen introduction.

The emergence of white-nose syndrome (WNS), caused by the introduced fungal pathogen, *P. destructans*^68–71^, has caused the collapse of North American bat populations^44,72,73^. Declines in three species (*Myotis lucifugus, Perimyotis subflavus,* and *Myotis septentrionalis*) have exceeded 90%^44,73–75^ and one species, *M. septentrionalis,* has been extirpated from most sites two to three years following pathogen arrival^76^. WNS is characterized by highly seasonal epizootics driven by interactions between thermal constraints of the pathogen (upper critical limit = 21°C), and the euthermic body temperature of bats (∼30°C)^77^. This limits active infections to the winter hibernation period when bats cool their body temperature to near ambient (1-15°C), for hibernation. If bats survive the winter, they can clear infections in the spring and mostly remain uninfected over the active summer season^77–79^. However, they become reinfected in the fall when they return to hibernacula and encounter contaminated environments and infected bats^77,80–82^.

Transmission of the multi-host pathogen, *P. destructans* is driven by both direct (host-to-host) and indirect (environment-to-host) contacts among hosts^73,80–82^. Previous work has demonstrated that cross-species and indirect transmission are important sources of infection needed to explain the rapid and widespread infections that have been observed in this system^81^. Declines during the epizootic have also been shown to be density dependent across multiple species^44,83^, suggesting that transmission is likely reduced in smaller colonies. Given the widespread declines observed from WNS, high cost of infection (mortality) and added costs of living in larger social groups (density-dependent declines), there should be considerable pressure to reduce interactions and limit transmission. Examination of species contributions to the environment pathogen reservoir (measured as propagule pressure) revealed that disease-induced declines did limit the importance of certain species following the establishment of WNS in sites^84^. Although other species increased in their importance following pathogen establishment, leaving an important question of whether transmission dynamics have changed over time^84^.

Determining whether transmission following disease establishment has changed and why has far-reaching implications for host adaptation, management and control, and recovery in highly impacted species.

Here, we use a surrogate pathogen (ultraviolet fluorescent powder) to simulate 134 unique pathogen introduction events originating from three bat species at two time periods, before the invasion of *P. destructans* (pre-WNS) and five to six years following disease-induced declines (post-WNS). We then used surrogate pathogen transfer to estimate the probability of inter and intra-species transmission, variability in exposure, environmental contamination, and habitat use across eight sites pre-WNS and six sites post-WNS. Generally, we find that transmission rates and exposure have increased post-WNS across all species, as well as the amount of surrogate pathogen contamination each bat contributes to the environment. In addition, we see shifts in space use within sites, with bats now using a larger portion of the hibernaculum than pre-WNS. Combined, these results suggest that transmission efficiency^85^ has increased in bat communities following severe declines and subsequent population stabilization.

## Methods

Using a surrogate pathogen, we examined within and among-species transmission rates, environmental pathogen shedding, and space use in bat populations before and again after the arrival of the fungal pathogen, *P. destructans* (pre- and post-WNS). Prior to the arrival of *P. destructans* (winters 2013/14 and 2014/15), we estimated transmission rates in eight sites in Wisconsin and Michigan (Supplemental Table 1). *Pseudogymnoascus destructans* arrived at all sites within a two-year period (2015-2017), and disease-induced mortality led to species-specific declines ranging between 48.3-100%, on average among species (Supplemental Table 1; *Eptesicus fuscus*: 48.3%, *Myotis lucifugus*: 88.7%, *M. septentrionalis*: 100%, *Perimyotis subflavus*: 89.8%). five to six years after the original study (2019/20), when populations had begun to stabilize^44,57^, we repeated the surrogate pathogen tracing at four of the original sites that still had bats present. Unfortunately, due to WNS mortality, four of the eight sites had < 2 bats present or had been completely extirpated (Supplemental Table 1). To increase our sample size for *P. subflavus*, we added one new site that matched the size and composition of the previously examined sites for a total of five post-WNS sites.

To estimate transmission rates over winter, we used a surrogate pathogen, ultraviolet fluorescent powder (UVFP), which has been previously described (Ref ^81^) in this system. The UVFP is capable of being transmitted through both direct and indirect contact and mirrors the transmission pathways of the epidermal fungal pathogen that causes WNS. Previous experimental work (Ref^81^, Methods section) has shown that UVFP transfer is mostly limited to a single chain of transmission (origin to primary recipient) and generally does not reflect onward transmission to secondary or tertiary recipients. This is likely due to the electrostatic properties (cationic (positively charged) surface for adhesion to surfaces) of the pigments (UVFP) used, which adheres to the negatively charged mammalian skin. Given the greatly reduced amount of UVFP that is transferred during an interaction between an origin and recipient bat, the powder is less likely to then transmitted onward from one recipient to another but is transmitted from our origin to other recipients. By using multiple colors of UVFP within the same sites we can repeatedly estimate transmission from a single introduction in the same communities but from different individuals and species, providing repeat measures in the same sites. Bats in this region begin hibernating between September–October and leave hibernacula in April–May. We applied the UVFP in November and conducted resighting in March to capture the core hibernation period when transmission occurs (∼90-day period (range: 88, 93)). During the summer prior to the second transmission study, at sites where we previously used the surrogate pathogen, we used a combination of high-pressure water (wildland fire backpacks) and scrub brushes to remove all remaining UVFP from the environment to ensure no environment-to-host transmission from the previous study.

We define each index bat that had UVFP applied in November as the "origin bat” and all bats capable of receiving each color of UVFP as the “recipients” (all bats in the site minus the origin bat). We applied UVFP to a total of 143 unique individual origin bats across three species between the two experimental periods. There were nine instances (6% of the total origin bats) where an origin bat did not make any contacts (no dust resighted of that color) and was not resighted in the site. We assumed these bats did not stay in the site or were predated and they were removed from further analyses, leaving a total of 134 origin bats in the final dataset (origin *M. lucifugus*: pre N=42, post N=30; *M. septentrionalis*: pre N=46, post N=0; *P. subflavus*: pre N=10, post N=6). When possible, we prioritized selecting origin bats for the post-WNS study that had been banded in previous years and survived at least one winter with WNS (Supplemental Fig. 1) to compare the post-experiment in surviving individuals. We applied UVFP to 3-7 individuals per site (Supplemental Table 2) using identical techniques among experiments ^81^. We applied 1g of UVFP to each individual origin bat, covering the body, wings and tail membrane, but avoiding application directly to the face. UVFP application occurred over a disposable bag to ensure there was no cross-contamination among colors or UVFP transfer from the site of application. One of 7 colors (ECO-11 Aurora Pink, ECO-15 Blaze Orange, ECO- 13 Rocket Red, ECO-17 Saturn Yellow, ECO-18 Signal Green, ECO-19 Horizon Blue (Day-Glo Color); and DFSB-C0 Clear Blue (Risk Reactor)) was randomly assigned to each individual bat, and each color was only used once per site. Following surrogate pathogen application, bats were allowed to recover in a clean towel to shed off any excess UVFP and then returned near to where they were captured.

When we returned in March for UVFP resighting, we first examined all bats present in the site with both visible light and ultraviolet (UV) light (wavelength ∼390nm) to determine which colors of dust were present on each recipient bat. We recorded the bats band or applied a new band if not previously marked to keep track of individuals. We counted all bats in the site by species and collected an epidermal swab from the bats forearm and muzzle to determine infection status using a previously described standardized technique^77^. To test for the presence and quantity of *P. destructans* (Supplemental Table 3), samples were extracted using a Qiagen DNeasy Blood and Tissue Kit and tested using an established *P. destructans*-specific qPCR ^77,86^. We resighted all bats before they aroused from hibernation to avoid disturbance-induced inflation in transmission. We also systematically surveyed all environmental surfaces (walls and ceilings) with visible and UV lights to count and quantify all UVFP contamination made by the origin bats in the environment. We also recorded the color and distance measured from the entrance of each environmental contact, when site structure allowed (single passage), to look at within-site space use (four sites pre and three sites post-WNS).

## Analysis

To analyze our data, we fit a series of Bayesian generalized linear mixed-effects (GLMM) and multivariate models, which are summarized in Supplemental Table 4. Unless otherwise noted all models were fit in “*brms”*^87^ using four MCMC chains with 4,000 iterations each (1,000 warm-up). Weakly informative priors were specified to improve sampling performance and reduce divergent transitions. Generally, fixed effects were assigned normal priors (mean = 0, SD = 10), while the intercept and random effect were assigned Student’s t priors (df = 3, location = 0, scale = 2.5). The target acceptance rate was increased (adapt delta = 0.99) to improve sampling efficiency and reduce divergent transitions when necessary.

Convergence was assessed using trace plots, Gelman–Rubin statistics (R^<1.01 for all parameters), and effective sample sizes (ESS generally >2000), all of which indicated good chain mixing and adequate sampling of the posterior distribution. No divergent transitions were observed.

We determined successful transmission as the transfer of UVFP from one of our origin bats to a recipient, which indicates either direct or indirect particle transfer. We then modeled transmission success pre- and post-WNS using a Bayesian GLMM. Our response variable was the number of successful transmission events (number of bats resighted with each color of UVFP) out of the total number of trials, modeled using a binomial distribution with a logit link function. The number of trials represents the total population size of each species at each site that were resighted for UVFP (minus the origin bat). Population-level (fixed) effects included origin species (*M. lucifugus*, *M. septentrionalis*, and *P. subflavus*), an interaction with recipient species (*Eptesicus fuscus*, *M. lucifugus*, *M. septentrionalis*, and *P. subflavus*), and an additive effect for experiment (pre- and post-WNS) to allow transmission probabilities to vary among all origin– recipient species combinations. Site was included as a group-level (random intercept) effect to account for non-independence among observations collected from the same site. Posterior predictions were generated using population-level effects and transformed to the response scale to estimate transmission probabilities among species pairs. Estimates are reported as means of the posterior distribution with 95% credible intervals. Model comparisons of previous experiments (Ref^81^) found no support for including color as a group-level effect (i.e., it did not make meaningful changes in posterior estimates or improve model accuracy), and therefore it was not included in further analyses.

We estimated posterior differences in transmission probability between pre- and post-WNS periods using the fitted Bayesian GLMM described above. A factorial prediction grid including all combinations of origin species, recipient species, and period (pre- and post-WNS) was constructed, and population-level expected transmission probabilities were generated using posterior predictions from the population-level effects only. To quantify the change between pre- and post-WNS transmission, posterior draws of predicted transmission probabilities were extracted using the full posterior distribution and paired by iteration, origin–recipient combination, and sampling period. For each posterior draw, the difference in transmission probability (post-WNS − pre-WNS) was estimated, yielding a full posterior distribution of temporal change for each species pair. These distributions were summarized using posterior means and 95% credible intervals to describe the changes in transmission probability across species combinations.

We then modeled how exposure in recipient bats changed pre- and post-WNS using the number of colors (unique origin contacts) received by individual recipient bats. We used a Bayesian GLMM with a Poisson distribution and log link function. The response variable was the number of colors received per individual recipient bat, and models accounted for variation in the number of origin bats by including an offset term for the log of the total number of colors available in a site during an experiment (effort within each site pre- & post-WNS). Population-level effects included recipient species (*E. fuscus*, *M. lucifugus*, *M. septentrionalis*, and *P. subflavus*) and an interaction with experimental period (pre- and post-WNS), and we included a group-level effect for site. Population level effects were assigned weakly informative normal priors (mean = 0, SD = 2). To interpret the interactive effects, we computed the estimated marginal means (EMMs) on the response scale using posterior draws and used pairwise contrasts between pre- and post-WNS conditions to estimate multiplicative change within each species.

Displayed model estimates are calculated using seven origin colors (mean: 6.81) to account for the offset term. In addition, heterogeneity in recipient exposure was quantified using the Gini coefficient, calculated from the proportion of total transmission events received by each individual host. Gini coefficients were estimated separately for each recipient species and experimental period (pre-WNS and post-WNS) using the Gini() function in the R package “*ineq”*. Higher Gini coefficients (max=1) indicate that transmission is concentrated to a relatively few individuals, whereas lower values indicate a more even distribution of exposure among hosts (min=0).

To examine the effect of pathogen infection (pathogen load) on the extent of surrogate pathogen transmission, we used a Bayesian GLMM with a binomial distribution where our response variable was whether individuals in the population either received a given color of UVFP or not (1|0). We included recipient species (*E. fuscus* and *M. lucifugus*) interacting with pathogen load as a continuous population-level effect and site as a group-level effect. This was examined for just a single origin species, *M. lucifugus*, because we had smaller sample sizes and an insufficient number of infected *P. subflavus* for analysis.

To determine whether there were differences in pathogen shedding into the environment (measured using a surrogate pathogen environmental contacts) across species and WNS period, we fit a Bayesian multivariate generalized linear mixed model. The model jointly estimated two response variables: (i) total size of environmental contamination per individual using a Gaussian distribution and (ii) the number of environmental contacts (*n*) using a negative binomial distribution, each with species (*M. lucifugus* and *P. subflavus*) and an additive effect for WNS period as our population-level effects and a shared group-level effect for site. We used a multivariate structure to account for potential covariance between the size of environmental contamination and number of contacts, while allowing both outcomes to share information on site-level structure. This framework also allows direct assessment of whether variation in total response is independent of variation in frequency of environmental contacts. We also examined the effect of pathogen infection on the extent of environmental contamination. We used a Bayesian GLMM with a Gamma error distribution and a log link function, where our response variable was log_10_ size of UVFP contamination in the environment. We included recipient species (*M. lucifugus* and *E. fuscus*) interacting with pathogen load as a continuous population-level effect and individual ID as a group-level effect.

We estimated the core area of space-use based on the inner quantiles (measured as distance in linear sites with one passage) of environmental contacts of each color of UVFP from an individual origin bat. We then modeled the proportional space-use (fraction of the site used based on environmental contacts to standardize among different size sites) using a Bayesian GLMM with a beta distribution and a logit link, which included experimental period (pre- vs post-WNS) as a population-level effect. Distances, used to estimate space-use, were only collected at four sites pre- and three site post-WNS so we were unable to include site as a group-level effect. However, we compared models with and without site as a population-level effect using Leave-One-Out cross-validation (LOO). The comparison showed minimal difference between models (Expected Log Pointwise Predictive Density (ELPD) difference = −0.7 ± 1.6), indicating no meaningful improvement in predictive accuracy with the inclusion of site, and therefore we report the model excluding site as a fixed effect.

Finally, we quantified *P. destructans* infection dynamics during the initial epizootic (pre-WNS) and the year of the post-WNS surrogate pathogen tracing. Since *P. destructans* was not present during the pre-WNS contact tracing, we used data from the first year following *P. destructans* arrival, which has been shown to be more consistent among sites than the invasion year (first year of arrival) given the variable arrival times among sites during winter ^80,88^. We used a Bayesian hurdle lognormal mixed-effects model to account for the zero-inflated and right-skewed distribution of fungal loads. The response variable was individual fungal load, modeled with a lognormal distribution conditional on presence, and a separate Bernoulli process governing the probability of non-detection (zero vs. non-zero load). Both components were modeled simultaneously in a hurdle framework using a logit link for the hurdle (zero-inflation) process and an identity link for the lognormal mean structure. Population-level effects included sampling date, species (*E. fuscus*, *M. lucifugus*, *M. septentrionalis*, and *P. subflavus*), and WNS phase (pre vs. post-WNS). Site was included as a group-level effect in both the conditional and hurdle components to account for repeated sampling within hibernacula. This structure allowed inference on both changes in infection intensity among infected individuals and changes in detection probability. All analyses were performed using R core team (XXX) in RStudio (Version 2026.04.0+526).

## Results

Generally, we found that transmission, exposure, pathogen shedding into the environment, and space use within hibernacula have all increased following the invasion and establishment of WNS into bat communities. Transmission probability varied substantially among origin and recipient species combinations (Fig. 1; Supplemental Table 5), and there was statistical support for higher transmission in the post-WNS period (Fig. 1; pre- vs post-WNS β = 1.26, 95% credible intervals (CrI): 1.00–1.53), indicating a general increase in transmission within and among species (Fig. 1-2A; Supplemental Table 5).

**Figure 1:**
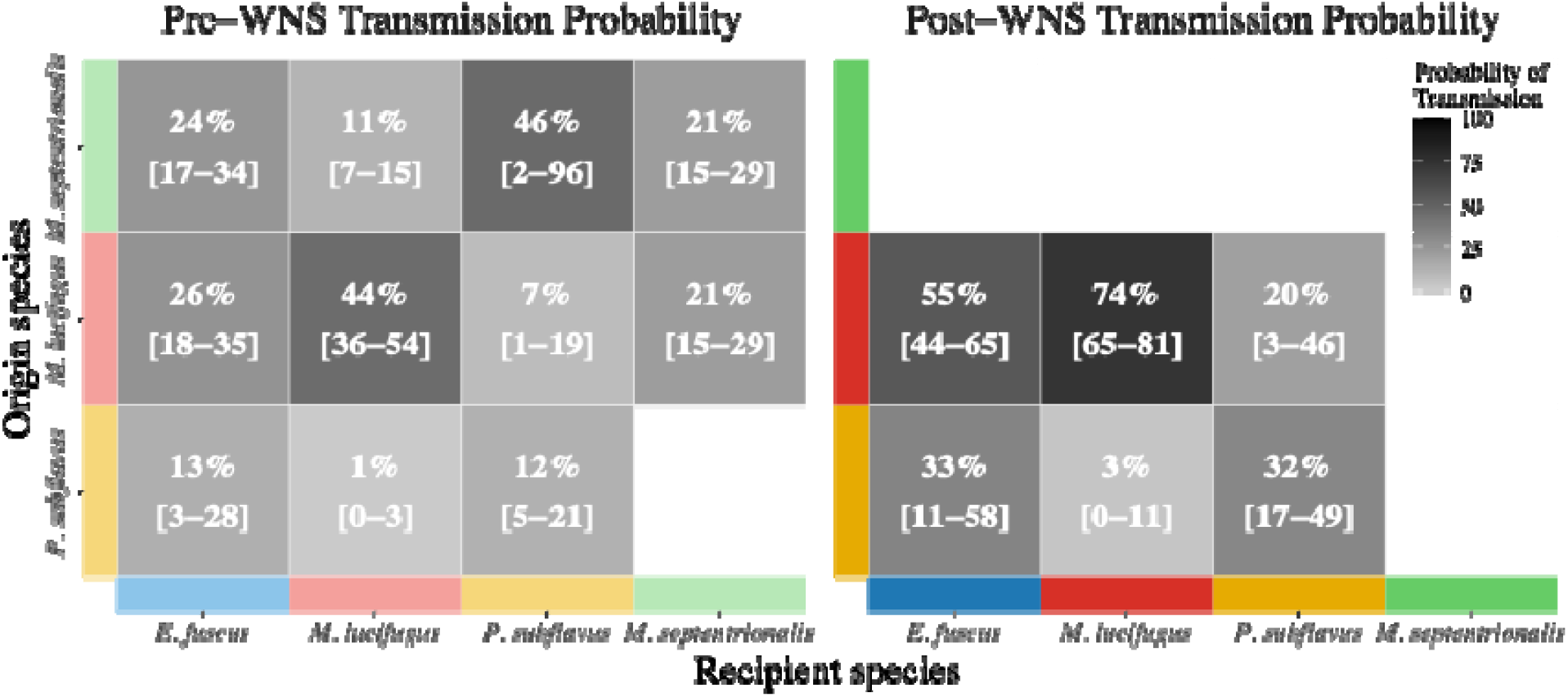
Inter- and intra-species transmission probabilities before and after WNS establishment. Heatmaps represent transmission probabilities originating from three species to all four species present in the community. Probability estimates shown within each tile represent the mean of the posterior distribution, and 95% posterior credible intervals are shown in brackets for each species pair. The response was modeled as binomial counts, and predictions were standardized to a fixed denominator of 100 trials per combination to facilitate direct comparison of transmission probabilities across groups. (Left panel) Transmission probabilities prior to the establishment of WNS (pre-WNS) as measured through successful transmission of a surrogate pathogen (UVFP). The missing tile (*P. subflavus: M. septentrionalis*) indicates insufficient data to estimate the coefficients between this species pair. (Right panel) Represents transmission probabilities 5-6 years after the establishment of WNS (post-WNS) in bat communities. *M. septentrionalis* was completely extirpated from all sites and so we were unable to estimate post-WNS transmission probabilities to or from this species.

**Figure 2:**
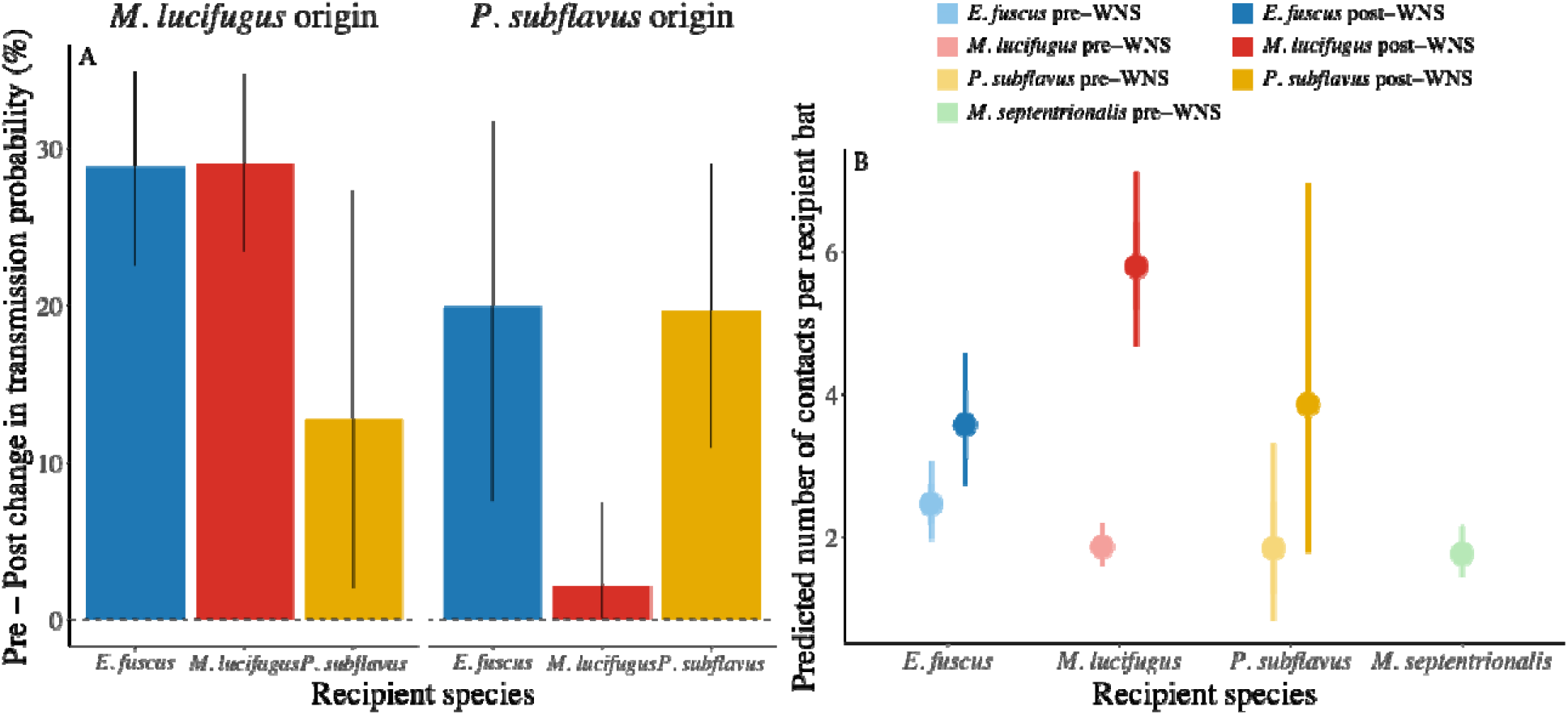
The change in transmission probability and exposure among species pre- to post- WNS establishment. (A) Posterior differences in transmission probability between pre- and post-WNS periods for *M. lucifugus* and *P. subflavus* origins. We generated population-level transmission probabilities using the full posterior distribution and paired by iteration. Bar plots show the posterior mean of the change and 95% CrI for each origin-recipient pair, and color indicates recipient species. (B) The panel shows the predicted number of origin bats that each recipient bat was in contact with (measured as the number of colors resighted on each recipient bat) during the pre- and post-WNS experiments. Points show the mean of the posterior distribution and whiskers show the 95% CrI (thin lines) and SD of the posterior (thick lines). The number of possible colors varied slightly among sites (mean: 6.81), which was included as an offset in the model. We used 7 colors in a site to predict the number of contacts to an individual bat pre- and post-WNS.

Before WNS establishment, *M. lucifugus* had higher transmission (Fig. 1 left panel) when serving as the source in comparison to *P. subflavus*, and to a lesser extent, *M. septentrionalis* (*P. subflavus*: β = −0.96, 95% CrI: -2.26 to 0.14; *M. septentrionalis* β = −0.08, 95% CrI: −0.52 to 0.37). Following the establishment of WNS in sites, transmission from *M. lucifugus* increased substantially (Fig. 2A, Supplemental Fig. 2, & Supplemental Table 6), with an increase in overall transmission probability across all recipient species (average 24% increase in transmission across two recipient species; Fig. 2A). Severe declines eliminated *M. septentrionalis* from the community (Table S1; 100% declines across all sites) eliminating their importance as a transmission source after WNS establishment. In the post-WNS experiment, we found support that the number of years a bat (*M. lucifugus*) survived with WNS (number of years banded) positively influenced transmission probability within the community (Supplemental Fig 3; Intercept = 0.18, 95% CrI: −0.84 to 1.22; years banded slope: β =0.33, 95% CrI: 0.09 to 0.57).

Exposure among recipient bats, measured as the number of colors each recipient received, also increased post-WNS, although effects varied among species (Fig 2B, Supplemental Table 7). *Myotis lucifugus* exhibited the strongest increase (Fig. 2B), rising from a predicted 1.53 (95% HPD: 1.28–1.78) colors per individual recipient pre-WNS to 4.77 (HPD: 3.84–5.84) colors post-WNS, representing a 3.12-fold increase (HPD: 2.68–3.62). *Eptesicus fuscus* showed a moderate increase (1.45-fold change; HPD:1.07–1.88) from 2.02 (HPD: 1.59– 2.51) to 2.93 (HPD: 2.23–3.77) colors. *Perimyotis subflavus* also increased from 1.45 (HPD: 0.59–2.61) to 3.05 (HPD: 1.24–5.43), although with less certainty (2.11-fold; HPD: 0.51–4.83). Due to declines in *Myotis septentrionalis* there was no data post-WNS for comparison (Fig. 2B). To assess whether exposure was evenly distributed among recipient individuals, we calculated the Gini coefficients (0=homogenous and 1= heterogenous) on the proportion of available colors received. Heterogeneity in surrogate pathogen exposure varied among species and between WNS periods, with evidence for reduced heterogeneity post-WNS in several taxa. *Myotis lucifugus* exhibited a marked reduction in heterogeneity, with Gini values decreasing from 0.205 pre-WNS to 0.097 post-WNS, alongside a large increase in mean proportional exposure. *Perimyotis subflavus* also showed a decline in inequality (0.397 to 0.218) concurrent with increased mean acquisition, although post-WNS estimates were based on smaller sample sizes. In contrast, *Eptesicus fuscus*, heterogeneity remained stable across periods (Gini 0.19 pre- and 0.20 post-WNS), despite an increase in mean exposure rates.

Origin bats with higher levels of infection (measured as quantities of pathogen DNA) showed increased cross-species transfer of UVFP (Fig. 3, Supplemental Table 8; *E. fuscus* β = 0.74, 95% CrI: 0.31 to 1.17). However, we found no support that transmission within species (other *M. lucifugus*) was influenced by infection severity (Fig. 3; *M. lucifugus* β = −0.21, 95% CrI: −0.63 to 0.19).

**Figure 3:**
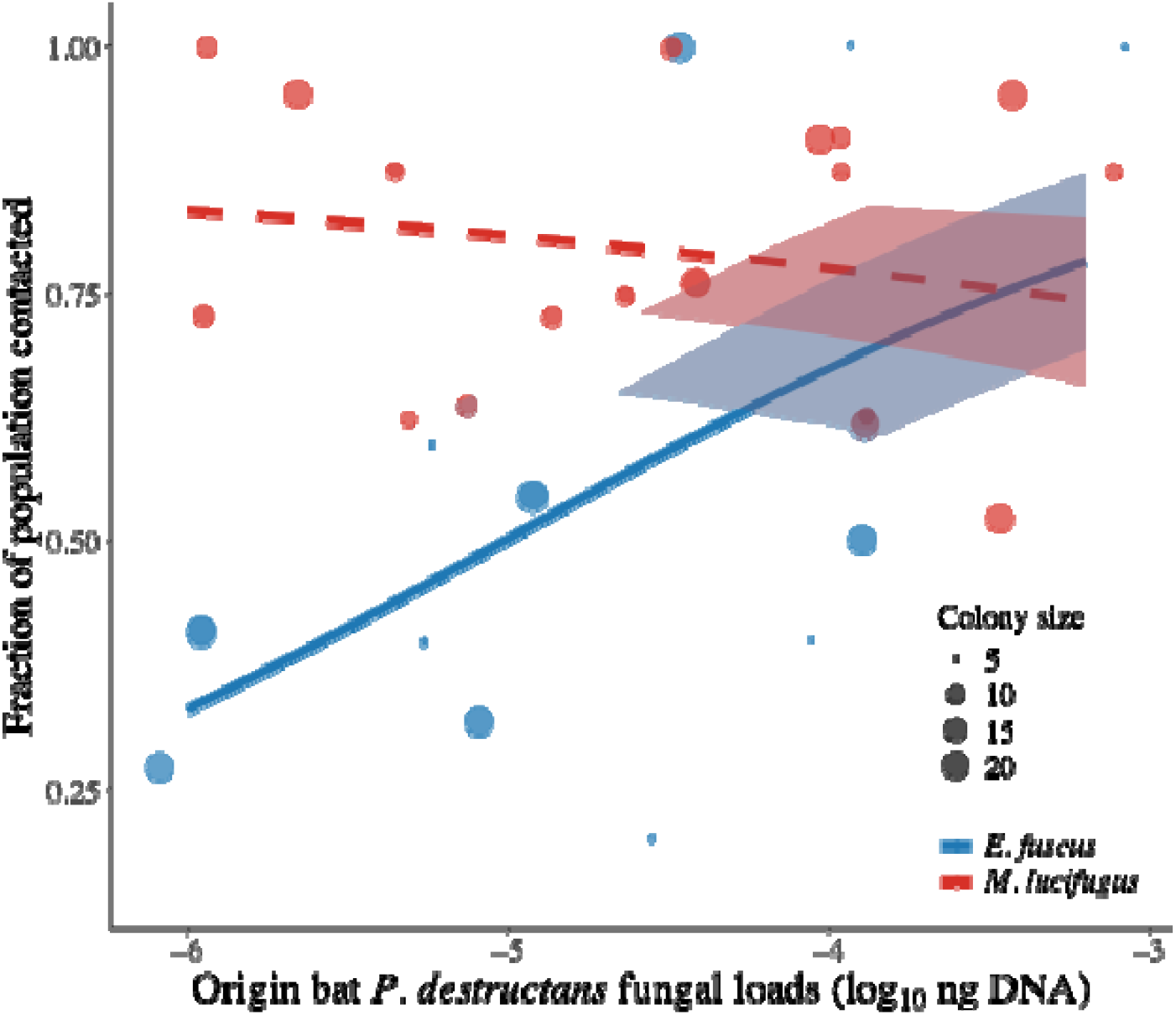
Effect of infection on inter and intra-species transmission. Points indicate the log_10_ population size for each species that was contacted by *M. lucifugus*. The lines indicate model fitted posterior means and line type indicates statistical support (dashed = CrI overlapping zero, solid = CrI not overlapping). On the x-axis is log_10_ fungal loads on *M. lucifugus* that were dusted in November and then recaptured at the end of the experiment (March). The y-axis is the fraction of each species in the site that was contacted and received the surrogate pathogen from the origin bat. Darker ribbons show the standard deviation of the posterior distribution and lighter ribbons show the 95% CrIs. There was a positive relationship between infection severity of the origin bat and the fraction of the *E. fuscus* population contacted (inter-species contacts β = 0.74, 95% CrI: 0.31 to 1.17) and no support for infection severity increasing the fraction of within-species contact for *M. lucifugus* (intra-species contact β = −0.21, 95% CrI: −0.63 to 0.19)

Examination of surrogate pathogen shedding into the environment showed that the amount of contamination in the environment (measured as total surface area (cm^2^) of UVFP resighted in the environment) was higher in post-WNS compared to the pre-WNS period for both *M. lucifugus* and *P. subflavus* (Fig. 4A, Supplemental Table 9). After WNS establishment, *M. lucifugus* had approximately a 73% increase in the surface area that an individual origin bat contaminated (Fig 4A; post-WNS β = 0.55, 95% CrI: 0.22–0.88). *Perimyotis subflavus* generally exhibited lower contamination than *M. lucifugus* (*P. subflavus* β = −0.43, 95% CrI: −0.75 to −0.08). There was also a general increase in the total number of environmental contacts following WNS emergence (Supplemental Table 9; β = 0.37, 95% CrI: −0.11 to 0.84); however, the credible interval overlapped zero, indicating weak support for a clear effect post-disease on contact frequency. Interestingly, we found no support that the amount of environmental contamination was influenced by infection intensity (Supplemental Fig. 4 & Supplemental Table 10; pathogen load, β = 0.12, 95% CrI: −0.14–0.39).

**Figure 4:**
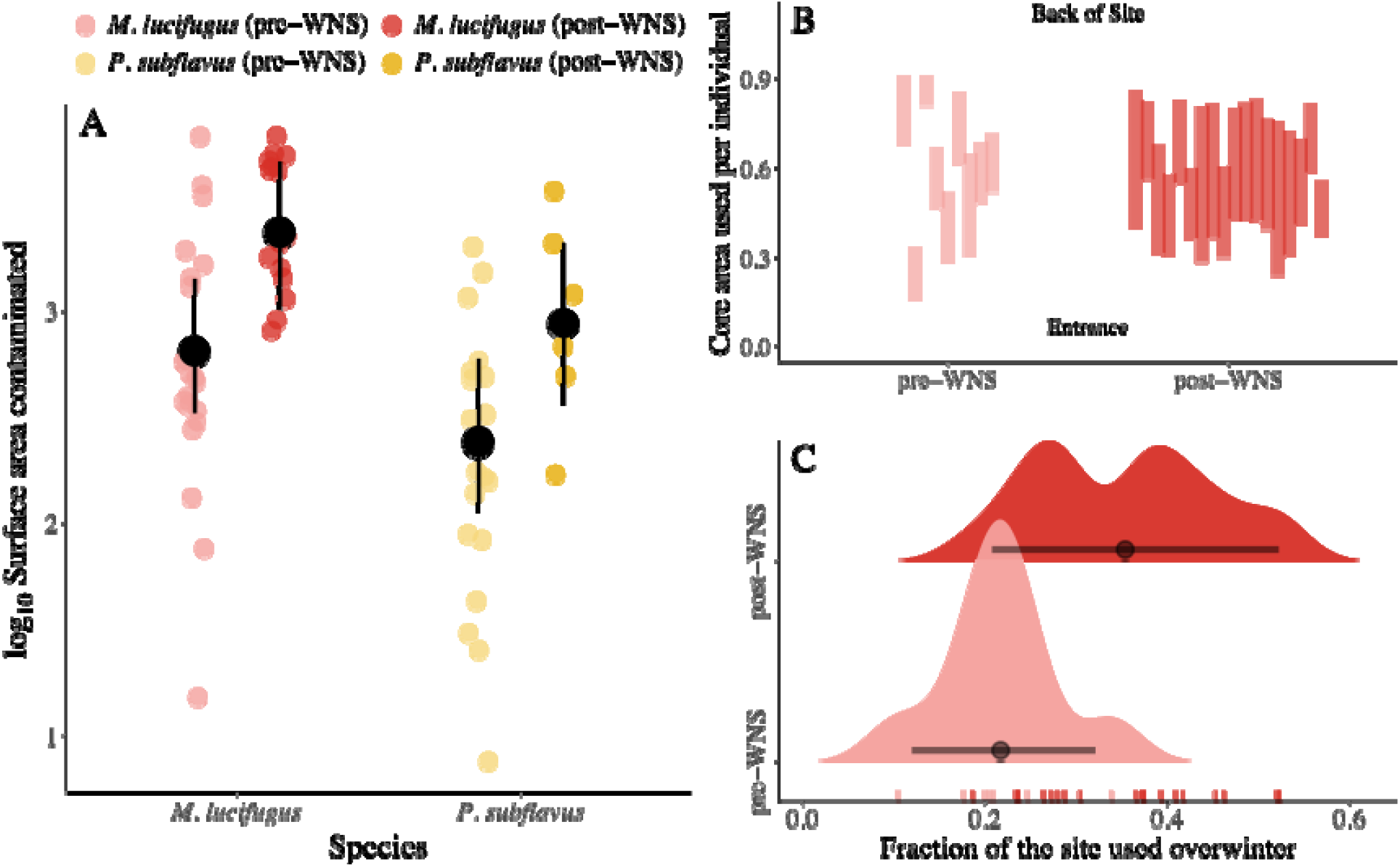
Changes in site use and environmental contamination following disease establishment. (A) Points indicate log_10_ total surface area (m^2^) of the environment contaminated with surrogate pathogen (UVF-dust) per individual. Larger black points and whiskers show the means and 95% CrIs. The color of each point indicates the origin species of the environmental contamination (*M. lucifugus* = red, *P. subflavus*=yellow). Darker shades of each color indicate the experimental period (Before WNS arrival or After WNS arrival), with light shades indicating surrogate pathogen transmission data from before WNS declines (pre) and darker shades indicate surrogate pathogen transmission data following declines (post = >5 years after *P. destructans* invasion). (B) The core area of use by each individual *M. lucifugus* within sites. Sites distances are normalized for comparison and shown as a fraction of the total site length. The core area is calculated as the 50% quantiles for locations contacted in the environment. (C) Shows the fraction of each site that was used by each individual bat calculated as the distance between the 25^th^ and 75^th^ quantiles (e.g. the length between upper and lower values for each bar in panel B) the black point indicates the posterior mean, and the lines indicate the 95% CrI. Rug plots at the bottom show that data for space use for each origin bat used in the density plots.

Space use by *M. lucifugus* pre- and post-WNS within hibernacula showed that individual bats now use a higher fraction of the total site following disease-induced declines (Fig. 4B-C, Supplemental Table 11; β = 0.66, 95% CrI: 0.29–1.04), which has generally consolidated in the interior part of the site and is less modular than pre-WNS (Fig. 4B). Distances were not recorded for *P. subflavus* so examination of changes in their space use was not possible.

Finally, examination of *P. destructans* infection dynamics pre-WNS (measured just after pathogen invasion) or post-WNS periods found no evidence that prevalence or infection intensity changed between the period when surrogate pathogen transmission was estimated (Fig. S5).

Posterior credible intervals for WNS effects in both components of the Bayesian hurdle lognormal model spanned zero, indicating no detectable shift in either prevalence or infection intensity following WNS establishment (Table S12).

## Discussion

Pathogen introduction events have resulted in the loss of biodiversity across the globe^21,44,89–94^. Ecological and evolutionary understanding of pathogen invasions and the processes that take place after can increase the effectiveness of intervention measures, facilitate species recovery, and help to better prepare for future introduction events^23,64,95–98^. The introduction of the fungal pathogen causing white-nose syndrome has caused severe declines in bat populations across North America^44,74,76^. The strong selective sweeps imposed by WNS have put pressure on bats to rapidly adapt or face local and regional extirpation^57,99–101^. Examination of transmission before and after the establishment of WNS has revealed that different components of transmission have been amplified within remnant bat communities. Following WNS establishment, when populations have stabilized, we see per-contact transmission probability, exposure, environmental contamination, and space use shift across surviving assemblages.

Together, these results suggest that post-WNS communities operate under higher transmission efficiency, but within a restructured network shaped by reduced diversity and increased dependence on a subset of remaining hosts.

Animal behavior can directly influence pathogen transmission by either facilitating or preventing a host from acquiring infection ^3,51,61^. Infection often requires significant investment from the host and redirects resource allocation from reproduction and survival to immune defenses and recovery^102–105^. Following pathogen introduction and widespread declines in novel host communities, we would predict that behaviors that accelerate their risk of exposure and infection would not be favored, particularly given that pathogen avoidance is generally thought to be less costly compared to investment in immunological defenses^106,107^. However, we see multiple facets of transmission and host-to-host interactions increase following disease establishment. The apparent increase in transmission contrasts with expectations for a density-dependent pathogen that imposes severe mortality on its hosts and suggests that the benefits of this increase far outweigh the costs of infection.

Recent work examining the relationships between bat population size and recovery in summer colonies when females communally raise their young revealed positive density-dependence in colony recovery (larger colonies had higher survival through WNS)^83^. This is counter to what has been observed in these same species and populations during the winter (density-dependent declines)^44,83^. The positive effect of larger colonies in the summer is likely due to the thermoregulatory benefit for heterothermic bats while raising their altricial young^108,109^. The pre-adaptive thermoregulatory need is likely exacerbated by the effects of WNS, which requires females to invest in healing during the reproductive period with even smaller population sizes. Thus increasingly favoring more social bats to form larger colonies ^83^. We also found that individual bats that had longer apparent survival (number of years since banding) had higher transmission probability. This could either be driven by selection against individuals with fewer social interactions (and lower transmission) or differences in social structure for older bats (more central in the network). Combined, these results suggest that the benefit of sociality (possibly driven by benefits in the summer) may outweigh the cost of increased social interactions during the winter that lead to a higher probability of transmission (Fig. 1).

Disease symptoms are known to both increase and decrease transmission. In WNS, bats are known to exhibit an increase in arousal frequency, which should provide more opportunity for host-to-host interaction and result in more pathogen shedding in the environment during the otherwise sedentary hibernation period. We found that while infection severity increased cross-species transmission (Fig. 3), it had no detectable effect on within-species transmission. In addition, we found no evidence to support that bats with higher infections contaminated more of the environment. Previous work has shown that within four to five years after the introduction of *P. destructans,* arousal patterns in bats return to their pre-WNS frequency^110^, which is further supported by the fact that bats are better able to budget their fat reserves in relation to infection only a few years after initial exposure^111^. This suggests that disease symptoms (increases in arousal frequencies) alone are unlikely to be driving the increase in interactions during the winter.

Generally, we found that disease dynamics among the two periods when transmission was estimated did not differ in prevalence or infection intensity (Fig. S5, except for M. septentrionalis which was extirpated from all sites), despite the dramatic decreases in population density (Table S1, >90% for three species). Given that several species have shown evidence of density-dependent declines^44,83^, it is possible that increases in transmission probability post-WNS establishment have offset the reduction that would otherwise be observed at reduced densities.

Our results demonstrate that pathogen invasions can fundamentally reorganize transmission systems long after the period of greatest mortality has passed. Rather than returning to pre-invasion conditions, remnant bat communities exhibit altered patterns of host contact, environmental interaction, and space use that collectively increase transmission despite dramatically reduced population sizes. Consequently, transmission parameters (e.g. contact rates) estimated prior to disease emergence may not accurately describe endemic dynamics in recovering populations and have consequences for future pathogen introductions. Understanding how host communities reorganize following pathogen invasion will therefore be critical for predicting long-term disease persistence, facilitating population recovery, and designing effective conservation strategies for wildlife threatened by emerging infectious diseases.

## Acknowledgements

The research was funded by the joint NSF-NIH-NIFA Ecology and Evolution of Infectious Disease award DEB-1911853, NSF DEB-1115895, and USFWS (F17AP00591). We would like to acknowledge landowners for access to the site A. Kurta, W. Scullon, and B. Heeringa for assistance in the field.

## Author Contributions

JRH wrote the original draft of the manuscript and conceptualized the project. JRH, and KEL designed methodology; JRH, and KEL contributed to funding acquisition; JRH, JED, NAL, MJK, HMK, JAR, JPW, and KEL performed field work; JRH and KEL performed data curation and formal analysis. All authors contributed critically to draft revision.

## Data Statement

The code and data required to reproduce all figures in this manuscript have been deposited in Figshare and are available for review at https://figshare.com/s/f3652fb93975247fb01a. Upon manuscript acceptance, the repository will be made publicly available under DOI: **10.6084/m9.figshare.33146141** (currently under embargo).

