## Supplemental Information for "Amplified transmission in host communities following disease-induced declines"

**Table of contents:**

**Figures**

Figure S1  Approximate age structure of bats used in the post-WNS transmission study
Figure S2  Transmission probability and total contacts pre- and post-WNS

Figure S3  Relationship between years since banding and transmission probability in post WNS *M. lucifugus*

Figure S4  Relationship between origin bat fungal loads and environmental contamination

Figure S5  Temporal dynamics of *Pseudogymnoascus destructans* prevalence and infection load across pre- and post-WNS periods

**Tables**

Table S1  Species-specific population counts across pre- and post-WNS periods

Table S2  Fraction of each recipient species contacted by each origin species (color)

Table S3  Prevalence and mean fungal loads by species and experimental period

Table S4  Summary of statistical models, figures, and output

Table S5  Species-pair transmission probabilities pre- and post-WNS

Table S6  Total contact events between species pairs across periods

Table S7  Exposure (number of colors received) in recipient species across WNS periods

Table S8  Effect of pathogen load on transmission probability from *M. lucifugus*

Table S9  Environmental contamination by origin species and WNS period

Table S10 Effect of pathogen load on environmental contamination size post-WNS

Table S11 Changes in within-hibernaculum space use by *M. lucifugus* pre- and post-WNS

Table S12 Lognormal hurdle model of *P. destructans* load and detection across WNS periods

**Supplemental Figures:**

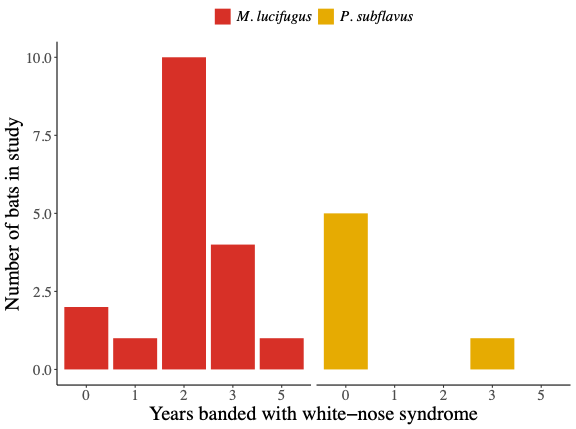

**Figure S1: Approximate age of bats used in the post-WNS transmission study.** We preferentially selected bats within each site that were banded in previous studies to better estimate transmission from surviving individuals. Bats were banded as part of a long-term project assessing survival and response to white-nose syndrome. All bats in the sites used in this study are banded every year. While bats cannot be reliably aged after their first summer, new individuals (unbanded bats that show up in sites each year) most likely represent juvenile recruitment to sites.

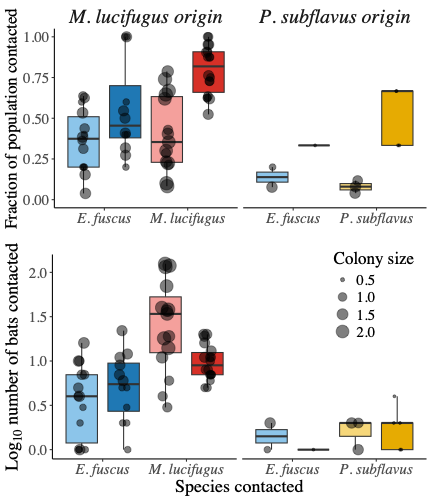

**Figure S2: Differences in transmission following disease-induced declines.** Points indicate the fraction of each species contacted by each origin bat (left *M. lucifugus* origin and right *P. subflavus* origin) and size of the point indicates log_10_ population for each species contacted. Boxplots show the lower (25^th^) and upper (75^th^) quartiles and whiskers show the range of data outside of the middle 50% of values. The color of each boxplox indicates the species that was contacted by *M. lucifugus* (left panel) or *P. subflavus* (right panel) (*E. fuscus* = blue, *M. lucifugus* = red, *P. subflavus*=yellow). Shades of each color indicate the period of the experiment. Light shades indicating pre-WNS surrogate pathogen transmission data from before WNS declines and darker shades indicate surrogate pathogen transmission data following declines (>5 years after *P. destructans* invasion).

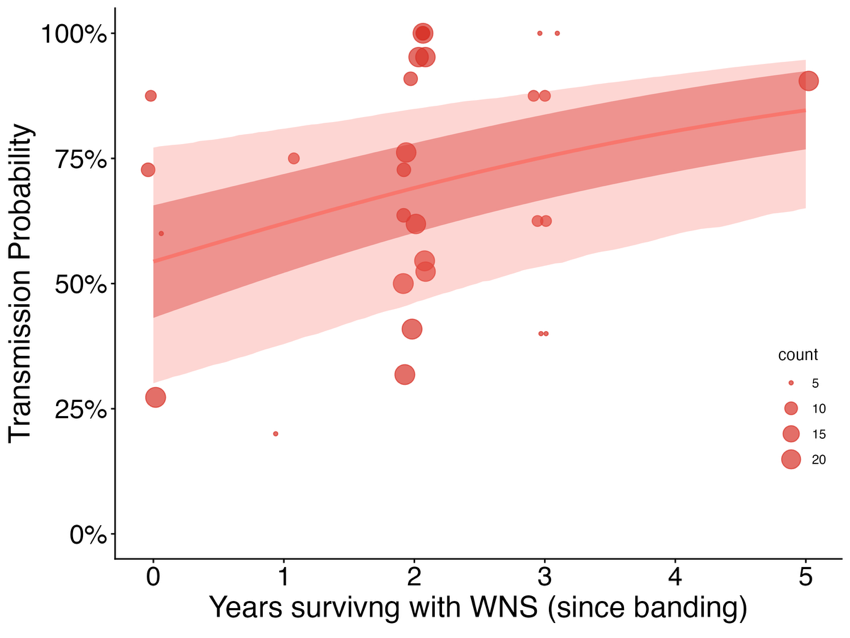

**Figure S3: Relationship between years since banding of the origin bat and surrogate pathogen transmission.** We estimated the effect of years since surviving WNS (years banded) of the origin bat on the probability of surrogate pathogen transmission by using a Bayesian GLMM with a binomial distribution, where UVFP contact sum | trials(count) ~ years since banding + (1 | site). This was restricted to just the post-WNS period and to *M. lucifugus*, which had enough variability in years since banding to merit examination. We found statistical support for a relationship (Intercept = 0.18, 95% CrI: -0.84 to 1.22; years banded slope: β =0.33, 95% CrI: 0.09 to 0.57) suggesting that bats that have survived longer have higher transmission probability.

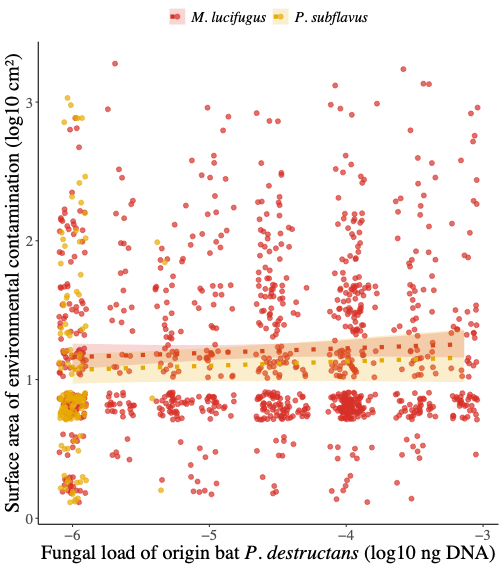

**Figure S4: Relationship between fungal loads on origin bats and the surface area they contaminated with UFVP.** Each point represents a surface-area measurement for an individual environmental contact and the fungal load of the origin bat that made the contact. Lines show the posterior means and the 95% CrIs.

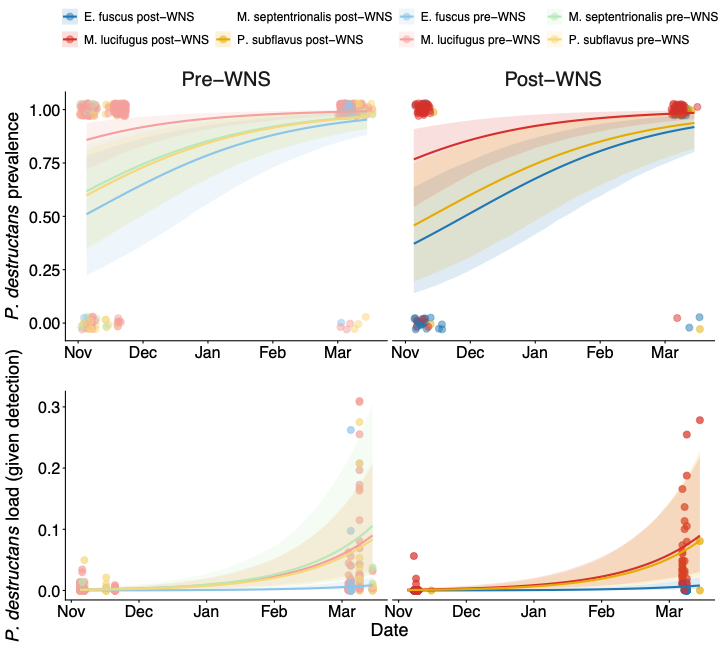

**Figure S5: Temporal dynamics of *Pseudogymnoascus destructans* infection prevalence and load across pre- and post-white-nose syndrome (WNS) periods.** The top panel shows model-estimated probability of *P. destructans* detection (prevalence), and the bottom panel shows predicted fungal load among infected individuals. Points represent raw observations (jittered for clarity). Lines indicate posterior mean predictions from Bayesian generalized linear mixed models, and shaded ribbons represent 95% credible intervals derived from posterior distributions. Pre-WNS data come from the first year following *P. destructans* arrival. *Pseudogymnoascus destructans* was not present in most sites (only two sites with late winter arrival) during the pre-WNS experiment. In addition, *P. destructans* arrival in the first year can be variable in sites occurring between fall and spring, and prevalence, loads, and mortality are low in the first year. Year two likely still represents the contact structure observed in the pre-WNS experiment but gives a more consistent look at changes in prevalence and loads over winter. Post-WNS data is from the same year (2019/20) that the post-experiment was conducted. *Myotis septentrionalis* was omitted from the post-WNS analysis since there were no individuals left to sample after WNS declines.

**Supplemental Tables:**

**Table S1: Population counts for each species by year.** Counts are shown for pre-WNS (2013/14 and 2014/15) and post-WNS (2019/20) in which transmission was estimated for sites. The percent decline is estimated between these two pairs of counts. White rows indicate the pre-WNS counts and gray rows indicate the post-WNS counts with % declines.

| **Site** | **Species** | **Winter year** | **Count** | **Population decline** |
| --- | --- | --- | --- | --- |
| BC | *E. fuscus* | 2014/15 | 16 |  |
| BC | *E. fuscus* | 2019/20 | 22 | 0% |
| BC | *M. lucifugus* | 2014/15 | 47 |  |
| BC | *M. lucifugus* | 2019/20 | 11 | 77% |
| BC | *M. septentrionalis* | 2014/15 | 19 |  |
| BC | *M. septentrionalis* | 2019/20 | 0 | 100% |
| BC | *P. subflavus* | 2014/15 | 2 |  |
| BC | *P. subflavus* | 2019/20 | 0 | 100% |
| GA | *E. fuscus* | 2014/15 | 30 |  |
| GA | *E. fuscus* | 2019/20 | 13 | 57% |
| GA | *M. lucifugus* | 2014/15 | 569 |  |
| GA | *M. lucifugus* | 2019/20 | 45 | 92% |
| GA | *M. septentrionalis* | 2014/15 | 39 |  |
| GA | *M. septentrionalis* | 2019/20 | 0 | 100% |
| HB | *E. fuscus* | 2019/20 | 1 | NA |
| HB | *P. subflavus* | 2019/20 | 3 | NA |
| MC | *E. fuscus* | 2014/15 | 11 |  |
| MC | *E. fuscus* | 2019/20 | 5 | 55% |
| MC | *M. lucifugus* | 2014/15 | 113 |  |
| MC | *M. lucifugus* | 2019/20 | 8 | 93% |
| MC | *M. septentrionalis* | 2014/15 | 19 |  |
| MC | *M. septentrionalis* | 2019/20 | 0 | 100% |
| SB | *E. fuscus* | 2013/14 | 5 |  |
| SB | *E. fuscus* | 2019/20 | 3 | 40% |
| SB | *M. lucifugus* | 2013/14 | 2 |  |
| SB | *M. lucifugus* | 2019/20 | 1 | 50% |
| SB | *M. septentrionalis* | 2013/14 | 1 |  |
| SB | *M. septentrionalis* | 2019/20 | 0 | 100% |
| SB | *P. subflavus* | 2013/14 | 10 |  |
| SB | *P. subflavus* | 2019/20 | 3 | 70% |
| SJ | *E. fuscus* | 2013/14 | 32 |  |
| SJ | *E. fuscus* | 2019/20 | 7 | 78% |
| SJ | *M. lucifugus* | 2013/14 | 205 |  |
| SJ | *M. lucifugus* | 2019/20 | 1 | 99.9% |
| SJ | *M. septentrionalis* | 2013/14 | 2 |  |
| SJ | *M. septentrionalis* | 2019/20 | 0 | 100% |
| SJ | *P. subflavus* | 2013/14 | 9 |  |
| SJ | *P. subflavus* | 2019/20 | 1 | 89% |
| SP | *E. fuscus* | 2013/14 | 5 |  |
| SP | *E. fuscus* | 2019/20 | 2 | 60% |
| SP | *M. lucifugus* | 2013/14 | 14 |  |
| SP | *M. lucifugus* | 2019/20 | 4 | 71% |
| SP | *M. septentrionalis* | 2013/14 | 5 |  |
| SP | *M. septentrionalis* | 2019/20 | 0 | 100% |
| SP | *P. subflavus* | 2013/14 | 25 |  |
| SP | *P. subflavus* | 2019/20 | 0 | 100% |
| ST | *M. lucifugus* | 2013/14 | 1.0 |  |
| ST | *M. lucifugus* | 2018 | 0.0 | 1.00 |
| ST | *M. septentrionalis* | 2013/14 | 1.0 |  |
| ST | *M. septentrionalis* | 2018 | 0.0 | 1.00 |
| ST | *P. subflavus* | 2013/14 | 17.0 |  |
| ST | *P. subflavus* | 2018 | 0.0 | 1.00 |
| TM | *M. lucifugus* | 2014/15 | 178 |  |
| TM | *M. lucifugus* | 2019/20 | 21 | 88% |
| TM | *M. septentrionalis* | 2014/15 | 66 |  |
| TM | *M. septentrionalis* | 2019/20 | 0 | 100% |

### Table S2: Fraction of each recipient contacted by each origin species and color in the pre-WNS and post-WNS experiment.

| **Site** | **Origin Species** | **Period** | **Winter** | **Recipient fraction *E. fuscus*** | **Recipient fraction *M. lucifugus*** | **Recipient fraction *P. subflavus*** | **Color** |
| --- | --- | --- | --- | --- | --- | --- | --- |
| BC | *M. lucifugus* | post-WNS | 2019/20 | 0.50 | 0.91 |  | b |
| BC | *M. lucifugus* | post-WNS | 2019/20 | 0.32 | 0.64 |  | g |
| BC | *M. lucifugus* | post-WNS | 2019/20 | 0.41 | 1.00 |  | o |
| BC | *M. lucifugus* | post-WNS | 2019/20 | 1.00 | 1.00 |  | r |
| BC | *M. lucifugus* | post-WNS | 2019/20 | 0.55 | 0.73 |  | w |
| BC | *M. lucifugus* | post-WNS | 2019/20 | 0.27 | 0.73 |  | y |
| BC | *M. lucifugus* | pre-WNS | 2014/15 | 0.44 | 0.23 |  | b |
| BC | *M. lucifugus* | pre-WNS | 2014/15 | 0.62 | 0.79 |  | o |
| BC | *M. lucifugus* | pre-WNS | 2014/15 | 0.31 | 0.40 |  | w |
| GA | *M. lucifugus* | pre-WNS | 2014/15 | 0.53 |  |  | l |
| HB | *P. subflavus* | post-WNS | 2019/20 |  |  | 0.67 | g |
| HB | *P. subflavus* | post-WNS | 2019/20 |  |  | 0.67 | o |
| HB | *P. subflavus* | post-WNS | 2019/20 |  |  | 0.33 | r |
| HB | *P. subflavus* | post-WNS | 2019/20 |  |  | 1.33 | w |
| MC | *M. lucifugus* | post-WNS | 2019/20 | 0.20 | 0.75 |  | b |
| MC | *M. lucifugus* | post-WNS | 2019/20 | 0.40 | 0.62 |  | g |
| MC | *M. lucifugus* | post-WNS | 2019/20 | 1.00 | 0.88 |  | o |
| MC | *M. lucifugus* | post-WNS | 2019/20 | 1.00 | 0.88 |  | r |
| MC | *M. lucifugus* | post-WNS | 2019/20 | 0.60 | 0.88 |  | w |
| MC | *M. lucifugus* | post-WNS | 2019/20 | 0.40 | 0.62 |  | y |
| MC | *M. lucifugus* | pre-WNS | 2014/15 |  | 0.35 |  | g |
| MC | *M. lucifugus* | pre-WNS | 2014/15 | 0.36 | 0.30 |  | p |
| MC | *M. lucifugus* | pre-WNS | 2014/15 | 0.64 | 0.62 |  | y |
| SB | *P. subflavus* | post-WNS | 2019/20 | 0.33 |  |  | g |
| SB | *P. subflavus* | post-WNS | 2019/20 |  |  | 0.33 | r |
| SB | *M. lucifugus* | pre-WNS | 2013/14 | 0.20 |  |  | l |
| SB | *M. lucifugus* | pre-WNS | 2013/14 | 0.60 |  | 0.10 | o |
| SB | *M. lucifugus* | pre-WNS | 2013/14 | 0.20 |  |  | y |
| SJ | *M. lucifugus* | pre-WNS | 2013/14 | 0.38 | 0.74 |  | l |
| SJ | *M. lucifugus* | pre-WNS | 2013/14 | 0.04 | 0.08 |  | o |
| SJ | *M. lucifugus* | pre-WNS | 2013/14 | 0.15 | 0.22 |  | y |
| SJ | *P. subflavus* | pre-WNS | 2013/14 |  | 0.01 |  | v |
| SJ | *P. subflavus* | pre-WNS | 2013/14 | 0.08 | 0.01 |  | w |
| SP | *M. lucifugus* | pre-WNS | 2013/14 | 0.40 | 0.43 | 0.04 | l |
| SP | *M. lucifugus* | pre-WNS | 2013/14 | 0.20 | 0.21 |  | o |
| SP | *M. lucifugus* | pre-WNS | 2013/14 |  | 0.29 |  | w |
| SP | *P. subflavus* | pre-WNS | 2013/14 |  |  | 0.08 | b |
| SP | *P. subflavus* | pre-WNS | 2013/14 | 0.20 |  | 0.04 | y |
| ST | *P. subflavus* | pre-WNS | 2013/14 |  |  | 0.12 | y |
| TM | *M. lucifugus* | post-WNS | 2019/20 |  | 0.76 |  | b |
| TM | *M. lucifugus* | post-WNS | 2019/20 |  | 0.62 |  | g |
| TM | *M. lucifugus* | post-WNS | 2019/20 |  | 0.90 |  | o |
| TM | *M. lucifugus* | post-WNS | 2019/20 |  | 0.95 |  | r |
| TM | *M. lucifugus* | post-WNS | 2019/20 |  | 0.95 |  | w |
| TM | *M. lucifugus* | post-WNS | 2019/20 |  | 0.52 |  | y |
| TM | *M. lucifugus* | pre-WNS | 2014/15 |  | 0.10 |  | b |
| TM | *M. lucifugus* | pre-WNS | 2014/15 |  | 0.67 |  | l |
| TM | *M. lucifugus* | pre-WNS | 2014/15 |  | 0.65 |  | w |

### Table S3 – Prevalence and average loads for each species by experimental period (pre- and post-WNS). Dates indicate the sampling period that corresponds to each experiment (pre-WNS = 2013/14 & 2014/15 and post-WNS = 2019/20). Sample sizes (N) during each sampling event indicate the number of each species sampled during that trip.

| **Site** | **Winter** | **Species** | **Date** | **N** | **Prevalence** | **Mean pathogen loads** |
| --- | --- | --- | --- | --- | --- | --- |
| BC | 2019/20 | *E. fuscus* | 2019-11-07 | 17 | 0.12 | -5.72 |
| BC | 2019/20 | *E. fuscus* | 2020-03-08 | 22 | 1.00 | -3.89 |
| BC | 2019/20 | *M. lucifugus* | 2019-11-07 | 14 | 0.71 | -4.67 |
| BC | 2019/20 | *M. lucifugus* | 2020-03-08 | 11 | 0.91 | -1.80 |
| BC | 2014/15 | *E. fuscus* | 2014-11-29 |  |  |  |
| BC | 2014/15 | *E. fuscus* | 2015-02-25 | 10 | 0.10 | -4.58 |
| BC | 2014/15 | *M. lucifugus* | 2014-11-29 | 13 | 0.08 | -4.57 |
| BC | 2014/15 | *M. lucifugus* | 2015-02-25 | 11 | 1.00 | -4.05 |
| BC | 2014/15 | *M. septentrionalis* | 2014-11-29 | 14 |  | -5.37 |
| BC | 2014/15 | *M. septentrionalis* | 2015-02-25 | 11 | 1.00 | -3.18 |
| BC | 2014/15 | *P. subflavus* | 2014-11-29 |  |  |  |
| BC | 2014/15 | *P. subflavus* | 2015-02-25 | 2 | 0.00 |  |
| GA | 2014/15 | *E. fuscus* | 2014-11-28 |  |  |  |
| GA | 2014/15 | *E. fuscus* | 2015-02-24 | 8 | 0.75 | -4.84 |
| GA | 2014/15 | *M. lucifugus* | 2014-11-28 | 13 | 0.00 |  |
| GA | 2014/15 | *M. lucifugus* | 2015-02-24 | 13 | 0.92 | -3.64 |
| GA | 2014/15 | *M. septentrionalis* | 2014-11-28 | 13 | 0.00 |  |
| GA | 2014/15 | *M. septentrionalis* | 2015-02-24 | 16 | 0.69 | -4.03 |
| GA | 2019/20 | *E. fuscus* | 2019-11-09 | 8 | 0.50 | -5.31 |
| GA | 2019/20 | *E. fuscus* | 2020-03-07 | 11 | 1.00 | -3.11 |
| GA | 2019/20 | *M. lucifugus* | 2019-11-09 | 1 | 1.00 | -0.88 |
| GA | 2019/20 | *M. lucifugus* | 2020-03-07 | 44 | 0.98 | -2.15 |
| GA | 2019/20 | *M. septentrionalis* | 2019-11-09 | 1 | 1.00 | -1.52 |
| GA | 2019/20 | *M. septentrionalis* | 2020-03-07 |  |  |  |
| HB | 2019/20 | *E. fuscus* | 2019-11-14 | 3 | 0.67 | -5.49 |
| HB | 2019/20 | *E. fuscus* | 2020-03-11 | 1 | 0.00 |  |
| HB | 2019/20 | *P. subflavus* | 2019-11-14 | 4 | 0.00 |  |
| HB | 2019/20 | *P. subflavus* | 2020-03-11 | 3 | 0.67 | -0.92 |
| MC | 2019/20 | *E. fuscus* | 2019-11-08 | 4 | 0.50 | -4.74 |
| MC | 2019/20 | *E. fuscus* | 2020-03-07 | 5 | 1.00 | -3.60 |
| MC | 2019/20 | *M. lucifugus* | 2019-11-08 | 10 | 1.00 | -3.96 |
| MC | 2019/20 | *M. lucifugus* | 2020-03-07 | 7 | 1.00 | -1.47 |
| MC | 2014/15 | *E. fuscus* | 2014-11-29 |  |  |  |
| MC | 2014/15 | *E. fuscus* | 2015-02-25 | 7 | 1.00 | -4.07 |
| MC | 2014/15 | *M. lucifugus* | 2014-11-29 | 13 | 0.08 | -4.70 |
| MC | 2014/15 | *M. lucifugus* | 2015-02-25 | 14 | 1.00 | -3.80 |
| MC | 2014/15 | *M. septentrionalis* | 2014-11-29 | 13 | 0.08 | -4.05 |
| MC | 2014/15 | *M. septentrionalis* | 2015-02-25 | 11 | 0.91 | -2.62 |
| SP | 2013/14 | *E. fuscus* | 2013-11-14 |  |  |  |
| SP | 2013/14 | *E. fuscus* | 2014-03-15 | 4 | 0.00 |  |
| SP | 2013/14 | *M. lucifugus* | 2013-11-14 | 8 | 0.00 |  |
| SP | 2013/14 | *M. lucifugus* | 2014-03-15 | 14 | 0.00 |  |
| SP | 2013/14 | *M. septentrionalis* | 2013-11-14 | 4 | 0.00 |  |
| SP | 2013/14 | *M. septentrionalis* | 2014-03-15 | 5 | 0.00 |  |
| SP | 2013/14 | *P. subflavus* | 2013-11-14 | 10 | 0.00 |  |
| SP | 2013/14 | *P. subflavus* | 2014-03-15 | 15 | 0.00 |  |
| SP | 2019/20 | *E. fuscus* | 2019-11-04 |  |  |  |
| SP | 2019/20 | *E. fuscus* | 2020-03-03 | 2 | 1.00 | -2.90 |
| SP | 2019/20 | *M. lucifugus* | 2019-11-04 | 2 | 1.00 | -3.50 |
| SP | 2019/20 | *M. lucifugus* | 2020-03-03 | 5 | 1.00 | -1.31 |
| SP | 2019/20 | *P. subflavus* | 2019-11-04 | 1 | 0.00 |  |
| SP | 2019/20 | *P. subflavus* | 2020-03-03 | 1 | 1.00 | -1.82 |
| SJ | 2013/14 | *E. fuscus* | 2013-11-09 |  |  |  |
| SJ | 2013/14 | *E. fuscus* | 2014-02-10 |  |  |  |
| SJ | 2013/14 | *E. fuscus* | 2014-03-21 | 10 | 0.00 |  |
| SJ | 2013/14 | *M. lucifugus* | 2013-11-09 | 9 | 0.00 |  |
| SJ | 2013/14 | *M. lucifugus* | 2014-02-10 |  |  |  |
| SJ | 2013/14 | *M. lucifugus* | 2014-03-21 | 20 | 0.00 |  |
| SJ | 2013/14 | *M. septentrionalis* | 2014-02-10 |  |  |  |
| SJ | 2013/14 | *M. septentrionalis* | 2014-03-21 | 1 | 0.00 |  |
| SJ | 2013/14 | *P. subflavus* | 2013-11-09 | 5 | 0.00 |  |
| SJ | 2013/14 | *P. subflavus* | 2014-02-10 |  |  |  |
| SJ | 2013/14 | *P. subflavus* | 2014-03-21 | 10 | 0.00 |  |
| SJ | 2019/20 | *E. fuscus* | 2020-02-28 | 4 | 1.00 | -3.86 |
| SJ | 2019/20 | *M. lucifugus* | 2020-02-28 | 1 | 1.00 | -0.63 |
| SJ | 2019/20 | *P. subflavus* | 2020-02-28 | 1 | 0.00 |  |
| ST | 2013/14 | *M. lucifugus* | 2013-11-09 |  |  |  |
| ST | 2013/14 | *M. lucifugus* | 2014-02-10 |  |  |  |
| ST | 2013/14 | *M. lucifugus* | 2014-03-20 | 1 | 0.00 |  |
| ST | 2013/14 | *M. septentrionalis* | 2014-02-10 |  |  |  |
| ST | 2013/14 | *M. septentrionalis* | 2014-03-20 | 1 | 0.00 |  |
| ST | 2013/14 | *P. subflavus* | 2013-11-09 | 9 | 0.00 |  |
| ST | 2013/14 | *P. subflavus* | 2014-02-10 |  |  |  |
| ST | 2013/14 | *P. subflavus* | 2014-03-20 | 16 | 0.00 |  |
| SB | 2013/14 | *E. fuscus* | 2014-03-20 | 5 | 0.00 |  |
| SB | 2013/14 | *M. lucifugus* | 2013-11-11 | 3 | 0.00 |  |
| SB | 2013/14 | *M. septentrionalis* | 2014-03-20 | 1 | 0.00 |  |
| SB | 2013/14 | *P. subflavus* | 2013-11-11 | 10 | 0.00 |  |
| SB | 2013/14 | *P. subflavus* | 2014-03-20 | 10 | 0.00 |  |
| SB | 2019/20 | *E. fuscus* | 2019-11-15 | 4 | 0.00 |  |
| SB | 2019/20 | *E. fuscus* | 2020-03-14 | 2 | 0.00 |  |
| SB | 2019/20 | *M. lucifugus* | 2020-03-14 | 1 | 1.00 | -0.56 |
| SB | 2019/20 | *P. subflavus* | 2019-11-15 | 2 | 0.50 | -5.54 |
| SB | 2019/20 | *P. subflavus* | 2020-03-14 | 3 | 0.67 | -3.31 |
| TM | 2019/20 | *M. lucifugus* | 2019-11-09 | 23 | 0.91 | -4.37 |
| TM | 2019/20 | *M. lucifugus* | 2020-03-06 | 21 | 1.00 | -1.51 |
| TM | 2014/15 | *M. lucifugus* | 2014-11-28 | 14 | 0.00 |  |
| TM | 2014/15 | *M. lucifugus* | 2015-02-24 | 15 | 0.07 | -5.56 |
| TM | 2014/15 | *M. septentrionalis* | 2014-11-28 | 14 | 0.00 |  |
| TM | 2014/15 | *M. septentrionalis* | 2015-02-24 | 15 | 0.00 |  |

**Table S4: Statistical analysis, figure and model output summary table.** Analyses are shown in the order they appear in the paper allowing with figure and tables.

| # | Model | Distribution | Figure | Table |
| --- | --- | --- | --- | --- |
| 1 | UVFP contact sum \| trials(count) ~ origin species * recipient species + period + (1 \| site) | Binomial | Fig. 1 | S5 |
| 2 | UVFP contact sum ~ origin + recipient * period + (1 \| site) | Negbinomial | Fig. S2 | S6 |
| 3 | UVFP contact sum \| trials(count) ~ Years banded + (1 \| site) | Binomial | Fig. S3 | In legend |
| 4 | Extracted posterior draws from model #1 | Binomial | Fig. 2A |  |
| 5 | # UVFP colors received ~ recipient species * period + offset(log(total colors available)) + (1 \| site), | Poisson | Fig. 2B | S7 |
| 6 | UVFP contact sum \| trials(count)~ pathogen load * recipient + (1 \| site) | Binomial | Fig. 3 | S8 |
| 7 | Total size of UVFP environmental contact~ origin species * period + (1 \| site); N of UVFP environmental contact ~ origin species* period + (1 \| site) | Gaussian, Negbinomial | Fig. 4A,  No Figure | S9 |
| 8 | log10(size of UVFP environmental contact) ~ pathogen load + origin species+ (1 \| individual ID) | Gamma | Fig. S4 | S10 |
| 9 | Inner quartiles of the site used (fraction) ~ period | Beta | Fig. 4B, C | S11 |
| 10 | Pathogen load ~ Date + Species + period + (1 \| site), hu ~ Date + Species + period + (1 \| site) | Hurdle:  Lognormal,  Bernoulli | Fig S5 | S12 |

### Table S5: Probability of transmission between pairs of species (origin: recipient) pre- and post-WNS establishment. Model output for Bayesian GLMM with UVFP contact sum | trials(count) ~ origin species * recipient species + period + (1 | site) and a binomial distribution. Results are graphically shown in Fig. 1. Estimates show the mean of the posterior distribution, Std Error indicates the standard deviation of the poster mean, and lower and upper limits indicate the 95% credible intervals (CrIs) given as the 2.5th (lower limit) and 97.5th (upper limit) percentiles of the posterior distribution.

| Term | Estimate | | | Std Error | | Lower limit | | Upper limit |
| --- | --- | --- | --- | --- | --- | --- | --- | --- |
| (Intercept: Origin(*M. lucifugus*): Recipient(*E. fuscus*) | | -0.901 | 0.209 | | -1.321 | | -0.476 | |
| Origin(*M. septentrionalis*) | | -0.385 | 0.216 | | -0.812 | | 0.036 | |
| Origin(*P. subflavus*) | | -1.067 | 0.600 | | -2.393 | | 0.011 | |
| Recipient(*M. lucifugus*) | | 0.707 | 0.141 | | 0.436 | | 0.986 | |
| Recipient(*M. septentrionalis*) | | -0.364 | 0.189 | | -0.737 | | 0.012 | |
| Recipient(*P. subflavus*) | | -1.938 | 0.845 | | -3.843 | | -0.533 | |
| Post-WNS | | 1.182 | 0.139 | | 0.908 | | 1.452 | |
| Origin(*M. septentrionalis*): Recipient(*M. lucifugus*) | | -1.552 | 0.231 | | -2.015 | | -1.104 | |
| Origin(*P. subflavus*): Recipient(*M. lucifugus*) | | -3.688 | 0.996 | | -5.789 | | -1.799 | |
| Origin(*M. septentrionalis*): Recipient(*M. septentrionalis*) | | 0.269 | 0.267 | | -0.240 | | 0.790 | |
| Origin(*P. subflavus*): Recipient(*M. septentrionalis*) | | 0.037 | 9.821 | | -19.399 | | 19.991 | |
| Origin(*M. septentrionalis*): Recipient(*P. subflavus*) | | 2.911 | 1.976 | | -1.084 | | 6.882 | |
| Origin(*P. subflavus):* Recipient(*P. subflavus*) | | 1.893 | 1.078 | | -0.126 | | 4.205 | |
| Site (SD of intercepts) | | 0.394 | 0.170 | | 0.181 | | 0.835 | |

### Table S6: Total number of contact events between pairs of species (origin: recipient) pre and post-WNS establishment. Model output for Bayesian GLMM with UVFP contact sum ~ origin + recipient * period + (1 | site) and negative binomial distribution, N=74. Results are shown graphically in Fig. S1. Estimates show the mean of the posterior distribution, Std Error indicates the standard deviation of the poster mean, and lower and upper limits indicate the 95% credible intervals (CrIs) given as the 2.5th (lower limit) and 97.5th (upper limit) percentiles of the posterior distribution.

| Term | Estimate | Std Error | Lower limit | Upper limit |
| --- | --- | --- | --- | --- |
| (Intercept Origin(*M. lucifugus*): Recipient(*E. fuscus*):pre-WNS) | 1.695 | 0.426 | 0.831 | 2.517 |
| Origin(*P. subflavus*) | -1.747 | 0.503 | -2.726 | -0.754 |
| Recipient(*M. lucifugus*) | 1.753 | 0.272 | 1.224 | 2.290 |
| Recipient(*P. subflavus*) | 0.475 | 0.653 | -0.811 | 1.757 |
| Post-WNS | 0.073 | 0.316 | -0.552 | 0.680 |
| Recipient(*M. lucifugus*)*:* post-WNS | -1.628 | 0.369 | -2.354 | -0.899 |
| Recipient(*P. subflavus*): post-WNS | 0.343 | 1.074 | -1.852 | 2.424 |
| Site (SD of intercepts) | 1.009 | 0.363 | 0.495 | 1.929 |

### Table S7: Exposure (number of colors received) in recipient species between pre- and post-WNS. Model output for Bayesian GLMM with number of UVFP colors received ~ recipient species * period + offset (log (total colors available)) + (1 | site), and a Poisson distribution. Results are graphically shown in Fig. 2B. Estimates show the mean of the posterior distribution, Std Error indicates the standard deviation of the posterior mean, and lower and upper limits indicate the 95% credible intervals (CrIs) given as the 2.5th (lower limit) and 97.5th (upper limit) percentiles of the posterior distribution.

| Term | Estimate | Std Error | Lower limit | Upper limit |
| --- | --- | --- | --- | --- |
| (Intercept – Recipient (*E. fuscus*, post-WNS) | -0.685 | 0.135 | -0.951 | -0.421 |
| Recipient(*M. lucifugus*) | 0.493 | 0.137 | 0.226 | 0.761 |
| Recipient(*M. septentrionalis*) | -0.305 | 7.133 | -14.235 | 13.728 |
| Recipient(*P. subflavus*) | 0.026 | 0.381 | -0.761 | 0.735 |
| Pre-WNS | -0.369 | 0.147 | -0.656 | -0.081 |
| Recipient(*M. lucifugus*):  Pre-WNS | -0.768 | 0.164 | -1.084 | -0.446 |
| Recipient(*M. septentrionalis*): Pre-WNS | -0.029 | 7.131 | -14.049 | 13.857 |
| Recipient(*P. subflavus*): Pre-WNS | -0.389 | 0.535 | -1.469 | 0.649 |
| Site (SD of intercepts) | 0.155 | 0.087 | 0.051 | 0.374 |

### Table S8: Effect of Pathogen loads on transmission probability originating from *M. lucifugus* to two recipient species post-WNS. Model output for Bayesian GLMM with UVFP contact sum | trials(count)~ pathogen load * recipient + (1 | site) with a binomial distribution, N=30. Results are shown graphically in Fig. 3. Estimates show the mean of the posterior distribution, Std Error indicates the standard deviation of the posterior mean, and lower and upper limits indicate the 95% credible intervals (CrIs) given as the 2.5th (lower limit) and 97.5th (upper limit) percentiles of the posterior distribution.

| Term | Estimate | Std Error | Lower limit | Upper limit |
| --- | --- | --- | --- | --- |
| (Intercept – Recipient(*M. lucifugus*)) | 0.375 | 1.048 | -1.691 | 2.348 |
| Pathogen load | -0.217 | 0.208 | -0.629 | 0.177 |
| Recipient(*E. fuscus*) | 3.319 | 1.421 | 0.552 | 6.136 |
| Pathogen load: Recipient(*E. fuscus*) | 0.960 | 0.294 | 0.388 | 1.548 |
| Site (SD of intercepts) | 0.441 | 0.625 | 0.009 | 2.087 |

### Table S9: Total environmental contamination by origin species and experimental period (pre- and post-WNS). Output for a Bayesian Hierarchical multivariate model with Total size of UVFP environmental contact~ origin species * period + (1 | site); and number (N) of UVFP environmental contact ~ origin species* period + (1 | site) with a Gaussian and Poisson distribution, respectively. Results are shown graphically in Fig. 4A. Estimates show the mean of the posterior distribution, Std Error indicates the standard deviation of the posterior mean, and lower and upper limits indicate the 95% credible intervals (CrIs) given as the 2.5th (lower limit) and 97.5th (upper limit) percentiles of the posterior distribution.

| Response | Term | Estimate | Std Error | Lower limit | Upper limit |
| --- | --- | --- | --- | --- | --- |
| Sum | (Intercept – Origin(*M. lucifugus*): pre-WNS) | 1,183.844 | 337.800 | 527.335 | 1,856.744 |
| N | (Intercept – Origin(*M. lucifugus*): pre-WNS) | 33.213 | 6.449 | 20.637 | 45.859 |
| Sum | Origin(*P. subflavus*) | -781.572 | 467.937 | -1,705.566 | 138.596 |
| Sum | Post-WNS | 1,849.070 | 474.008 | 913.601 | 2,788.543 |
| Sum | Origin(*P. subflavus*): post-WNS | -954.528 | 838.180 | -2,596.977 | 680.563 |
| N | Origin(*P. subflavus*) | -17.812 | 8.683 | -34.899 | -0.587 |
| N | Post-WNS | 20.798 | 8.495 | 3.928 | 37.469 |
| N | Origin(*P. subflavus*): post-WNS | -11.768 | 15.108 | -41.913 | 17.549 |
| Site | Site (SD of random intercepts - Sum) | 286.057 | 229.904 | 10.837 | 867.196 |
| Site | Site (SD of random intercepts - N) | 7.635 | 6.471 | 0.267 | 24.168 |
| Sum | Residual SD (Sum) | 1,430.713 | 135.102 | 1,190.221 | 1,722.370 |
| N | Residual SD (N) | 24.980 | 2.555 | 20.292 | 30.350 |

### Table S10: Effect of pathogen infection on size of environmental contamination post-WNS. Model output for a Bayesian GLMM where log10(size of UVFP environmental contact) ~ pathogen load + origin species+ (1 | individual ID) with a Gamma distribution. Results are shown graphically in Fig. S2. Estimates show the mean of the posterior distribution, Std Error indicates the standard deviation of the posterior mean, and lower and upper limits indicate the 95% credible intervals (CrIs) given as the 2.5th (lower limit) and 97.5th (upper limit) percentiles of the posterior distribution.

| Term | Estimate | Std Error | Lower limit | Upper limit |
| --- | --- | --- | --- | --- |
| (Intercept – Origin (*M. lucifugus*)) | 0.310 | 0.110 | 0.091 | 0.522 |
| Pathogen load | 0.027 | 0.024 | -0.021 | 0.073 |
| Origin(*P. subflavus*) | -0.083 | 0.064 | -0.206 | 0.045 |
| Individual ID (SD of intercepts) | 0.049 | 0.026 | 0.004 | 0.104 |

### Table S11: Change in within hibernacula space use by *M. lucifugus* pre- and post-WNS. Model output for a Bayesian GLMM where we compared the Inner quartiles of the site used (fraction) ~ period using a Beta distribution. Results are shown graphically in Fig. 4C. Estimates show the mean of the posterior distribution, Std Error indicates the standard deviation of the posterior mean, and lower and upper limits indicate the 95% credible intervals (CrIs) given as the 2.5th (lower limit) and 97.5th (upper limit) percentiles of the posterior distribution.

| Term | Estimate | Std Error | Lower limit | Upper limit |
| --- | --- | --- | --- | --- |
| (Intercept – pre-WNS) | -1.27 | 0.163 | -1.595 | -0.951 |
| Post-WNS | 0.66 | 0.190 | 0.288 | 1.040 |

### Table S12: Temporal dynamics of *Pseudogymnoascus destructans* infection prevalence and load across pre- and post-white-nose syndrome (WNS) periods. Model output for a Bayesian hurdle lognormal mixed-effects model with Pathogen load ~ Date + Species + period + (1 | site), hu ~ Date + Species + period + (1 | site), and a lognormal and Bernoulli distribution. Results are shown graphically in Fig. S5. Estimates show the mean of the posterior distribution, Std Error indicates the standard deviation of the posterior mean, and lower and upper limits indicate the 95% credible intervals (CrIs) given as the 2.5th (lower limit) and 97.5th (upper limit) percentiles of the posterior distribution.

| **Term** | **Estimate** | **Std Error** | **Lower limit** | **Upper limit** |
| --- | --- | --- | --- | --- |
| Intercept – (*E. fuscus*, post-WNS) | -21.65 | 0.96 | -23.55 | -19.76 |
| Hu Intercept – (*E. fuscus*, post-WNS) | 8.65 | 1.53 | 5.85 | 11.78 |
| Date | 0.92 | 0.05 | 0.82 | 1.03 |
| Species (*M. lucifugus*) | 2.36 | 0.33 | 1.72 | 3.01 |
| Species (*M. septentrionalis*) | 2.44 | 0.56 | 1.33 | 3.54 |
| Species (*P. subflavus*) | 2.28 | 0.49 | 1.31 | 3.22 |
| pre-WNS | 0.02 | 0.27 | -0.50 | 0.55 |
| Hurdle: Date | -0.73 | 0.11 | -0.94 | -0.53 |
| Hurdle: (*M. lucifugus*) | -1.83 | 0.47 | -2.76 | -0.94 |
| Hurdle: (*M. septentrionalis*) | -0.47 | 0.61 | -1.68 | 0.70 |
| Hurdle: (*P. subflavus*) | -0.38 | 0.58 | -1.52 | 0.73 |
| Hurdle: pre-WNS | -0.61 | 0.40 | -1.38 | 0.19 |
| Site (SD intercept, load model) | 1.04 | 0.41 | 0.50 | 2.04 |
| Site (SD intercept, hurdle model) | 1.03 | 0.51 | 0.37 | 2.36 |
| Residual SD (sigma) | 2.23 | 0.07 | 2.10 | 2.38 |
